# A Dual-Action Mechanism to Prevent OX40 Signaling: The Structural Basis for the Differentiated Antagonist STAR-0310

**DOI:** 10.1101/2025.09.16.676329

**Authors:** Nikolaos Biris, Chunxia Zhao, Julie Macoin, James R. Fuller, Andrew Blauvelt, Raj Chovatiya, Christopher G. Bunick

## Abstract

Atopic dermatitis (AD) is a chronic inflammatory disease sustained by dysregulated T cell activity. The OX40/OX40L pathway drives effector and memory T cell proliferation, survival, and cytokine production, making it a key therapeutic target. STAR-0310, a novel anti-OX40 antibody, binds a noncanonical epitope that sterically blocks receptor trimerization without inducing agonism. Structural and functional studies demonstrated a dual mechanism: prevention of new OX40/OX40L interactions and efficient disruption of pre-formed complexes, outperforming comparator antibodies. The pure antagonism and complex disruption capacity of STAR-0310 support its clinical evaluation (NCT06782477) as a differentiated OX40-targeted therapy for AD.

**Highlights:**

- **Novel binding mechanism:** STAR-0310 engages OX40 distal to the OX40L site, sterically blocking receptor trimerization without inducing agonism.
- **Dual action:** Prevents formation of new OX40/OX40L complexes and efficiently disrupts pre-formed complexes sustaining inflammation.
- **Differentiation from competitors:** Achieves greater efficiency in complex disruption compared with rocatinlimab and IMG-007 with no partial agonist activity.
- **Clinical potential:** Pure antagonist profile supports ongoing evaluation of STAR-0310 (NCT06782477) as a best-in-class OX40 therapy for atopic dermatitis.

---

Atopic dermatitis (AD) is a chronic, inflammatory skin disease with a complex, heterogeneous pathophysiology. Immunopathogenesis in AD is driven by T cell dysregulation that extends beyond the T helper (Th) 2 axis to include Th1, Th17, and Th22 T cell subsets^1,2,3^. Abnormally modulated T cell activity perpetuates a cycle of inflammation and skin barrier dysfunction, contributing to a significant quality-of-life burden for patients. This underscores the need for AD therapies that provide comprehensive and durable control over diverse inflammatory pathways^4,5^.

The co-stimulatory tumor necrosis factor receptor OX40 (CD134, TNFRSF4), expressed on activated T-cells, and its ligand, OX40L (CD252, TNFSF4), form a critical immunologic axis that acts as a key regulator of T cell activity. Engagement of this pathway promotes survival, proliferation, polarization and cytokine production of pro-inflammatory effector and memory T cell populations, while simultaneously inhibiting suppressive function of regulatory T cells^4,5^. Consequently, blockade of OX40/OX40L interactions has emerged as a promising therapeutic strategy for AD, and several biologics that target either OX40 or OX40L are currently in clinical development^6,7^.

However, the specific mechanism of antibody-mediated antagonism is a critical factor underlying therapeutic potential in this treatment class^8^. Here, the structural and functional basis for the differentiated mechanism of STAR-0310, a novel, investigational anti-OX40 antibody is described. STAR-0310 utilizes a unique binding mode that induces steric hindrance to achieve potent and pure antagonism which may translate to high efficacy in treating OX40-mediated inflammatory diseases.

To define the molecular basis of OX40 inhibition for STAR-0310 (and to compare it with other known anti-OX40 antibodies), high-resolution cryogenic Electron Microscopy (cryo-EM) structures were determined for STAR-0310, as well as rocatinlimab, and IMG-007 analogs (hereafter referred to as rocatinlimab and IMG-007 respectively), each in complex with human OX40^9^. The extracellular region of OX40 is composed of four cysteine-rich domains (CRD1-4), with the native OX40L trimer binding to an interface formed by CRD1 and CRD2^10^ (Figure 1A). Structural analysis revealed that STAR-0310 binds to an epitope opposite of the canonical OX40L binding interface (Figure 1B). This binding mode of STAR-0310 is distinct from that of rocatinlimab, which targets the CRD3 domain. Although rocatinlimab is engineered to deplete OX40⁺ T cells via enhanced antibody-dependent cellular cytotoxicity (ADCC), Fc-dependent clearance may not eliminate all targets^11^. Therefore, direct steric blockade by binding directly beneath the OX40 ligand-binding site at the CRD3 (Figure 1C) is important for inhibiting OX40 signaling on residual T cells. Although IMG-007 binds to an epitope that overlaps with that of STAR-0310 in CRD2, IMG-007 engages the receptor with a distinctly different fragment antigen binding (Fab) orientation (Figure 1D). The binding of STAR-0310 does not directly occlude the ligand docking interface but instead prevents the assembly of a signaling complex through potent steric hindrance. The trimeric nature of OX40L requires the recruitment and subsequent trimerization of three OX40 receptor monomers to initiate downstream nuclear factor kappa B (NF-κB) signaling. The binding of the STAR-0310 Fab region sterically obstructs the binding of the native OX40L trimer, thereby preventing the formation of the functional hexameric OX40/OX40L signaling complex (Figure 1E).

**Figure 1.**
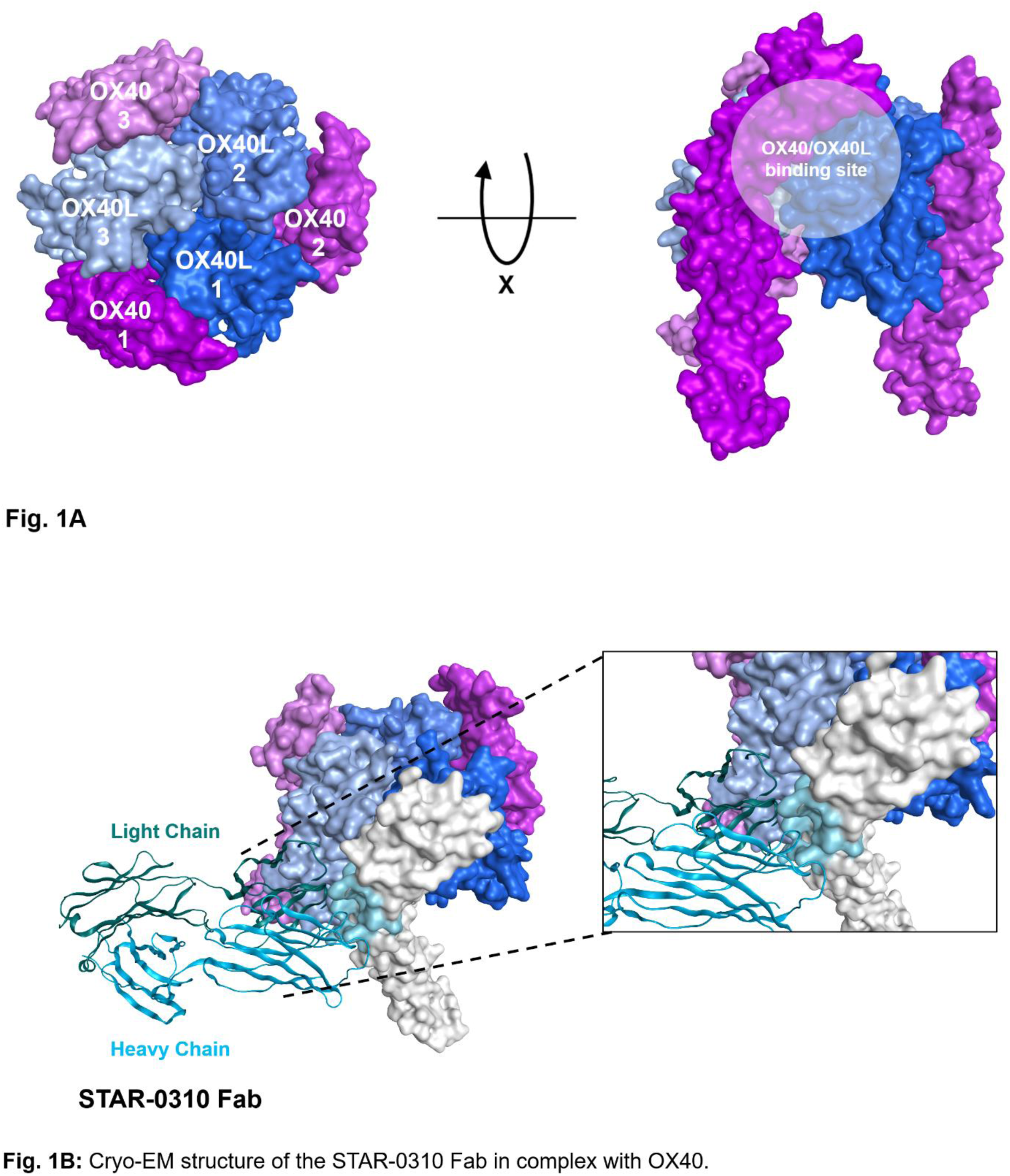
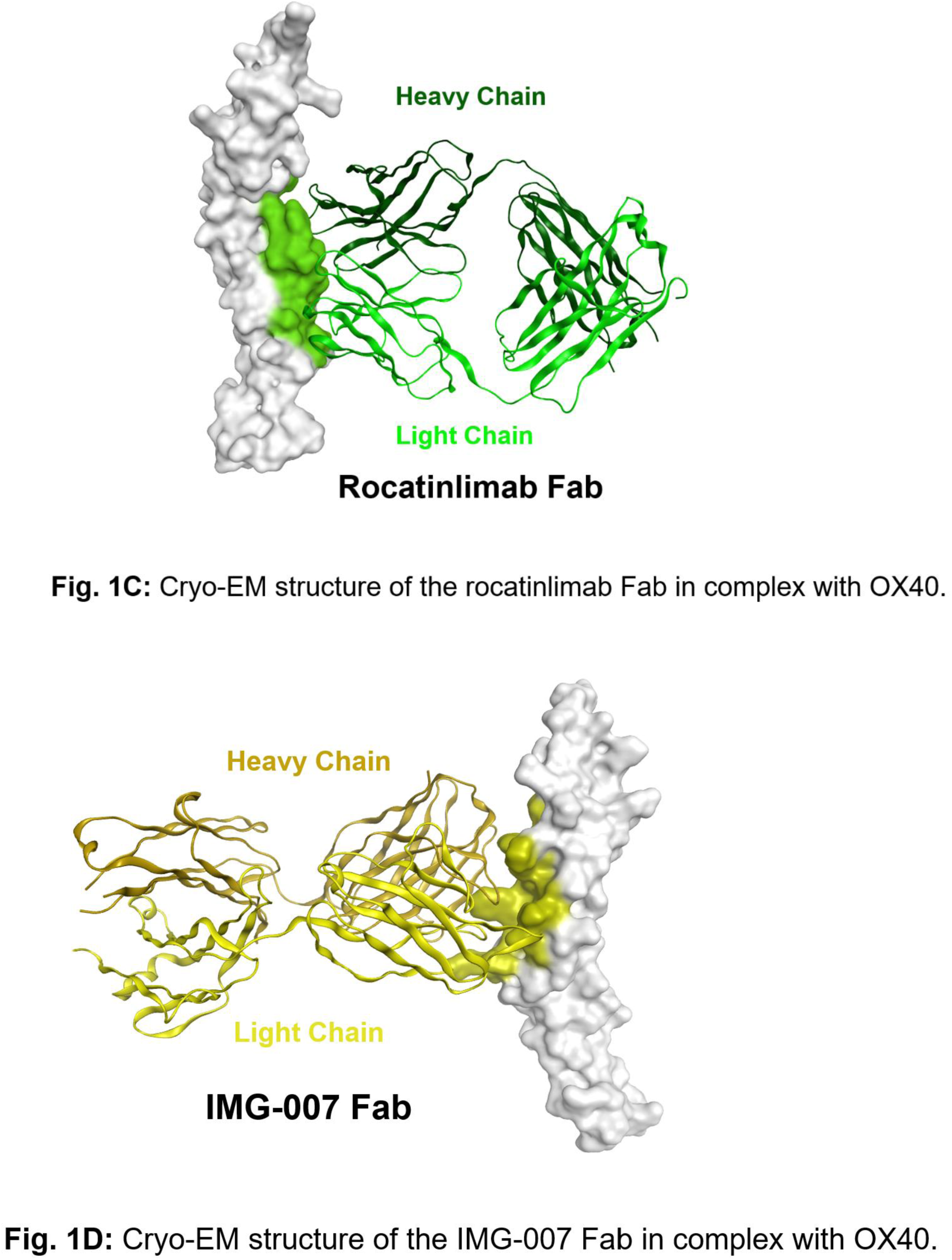
Structural basis for the differentiated mechanisms of anti-OX40 antibodies. (A) On the left, top-down surface view of the hexameric OX40:OX40L assembly (PDB 2HEV), comprising three monomeric OX40 molecules (purple) bound to an OX40L homotrimer (blue). On the right, side view of the same complex, obtained by a 90° rotation about the x-axis. The interaction interface is localized to Cysteine-Rich Domains (CRD) 1 and 2 of OX40. (B) Cryo-EM structure of the STAR-0310 Fab in complex with OX40. STAR-0310 binds a distinct epitope on CRD2 (light blue), at a site opposite the OX40L binding face. The STAR-0310 Fab/OX40 complex is depicted in the context of an OX40/OX40L hexamer. The inset provides a zoomed-in view of how the binding of STAR-0310 prevents the formation of the OX40/OX40L hexamer formation. (C) Cryo-EM structure of the rocatinlimab Fab in complex with OX40. Rocatinlimab binds to CRD3 (light green), directly beneath the OX40L binding site, where it acts via direct steric hindrance. (D) Cryo-EM structure of the IMG-007 Fab in complex with OX40. IMG-007 binds an epitope in CRD2 that overlaps with the STAR-0310 epitope (olive color) but displays a significantly different Fab orientation.

In chronic immune-mediated diseases like AD, pathogenic memory T cells are maintained by persistent signaling through established OX40/OX40L complexes, contributing to disease chronicity^12^. Therefore, an ideal therapeutic targeting OX40/OX40L should not only prevent the formation of new signaling OX40/OX40L complexes but also possess the ability to disrupt existing complexes to resolve established inflammation. A Surface Plasmon Resonance (SPR)-based complex disruption assay was utilized to assess this capability. Pre-formed OX40/OX40L complexes were immobilized on a sensor chip, and the ability of various antibodies to induce dissociation was measured in real-time. STAR-0310 induced a robust and rapid disruption of these pre-formed complexes, with a rate and efficiency approximately twice that observed for rocatinlimab and IMG-007 analogs (Figure 2A). By contrast, antibodies targeting OX40L, such as amlitelimab and APG990 in-house generated analogs (hereafter referred to as amlitelimab and APG990 respectively), were unable to disrupt the pre-formed complexes (Figure 2B), likely because their epitopes on OX40L are masked upon OX40L binding to OX40^13^. This potent ability to actively disrupt existing complexes is a key mechanistic differentiator for STAR-0310, which may potentially translate into more rapid and/or deeper clinical responses by targeting interactions that sustain inflammation in established AD lesions.

**Figure 2.**
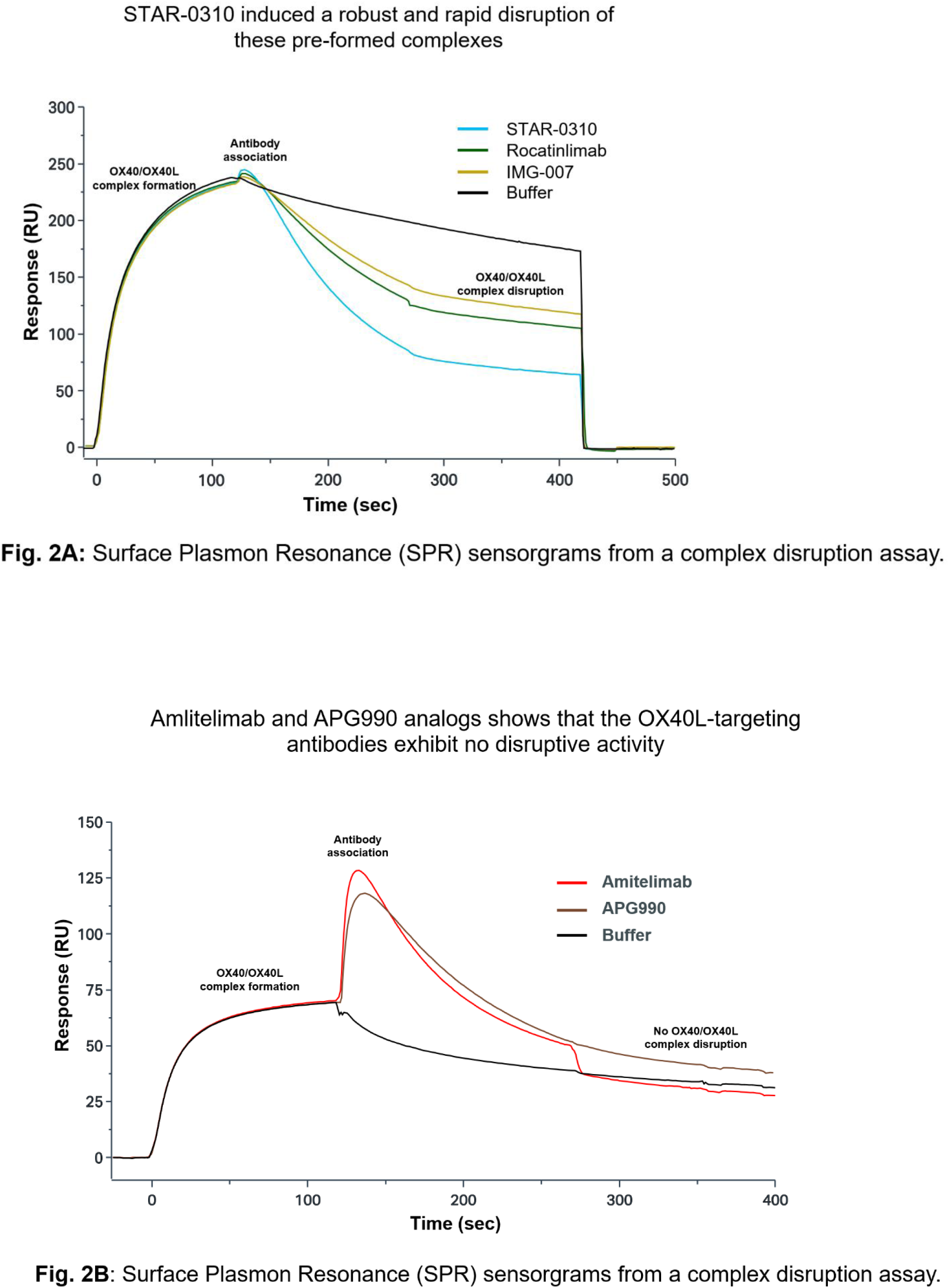
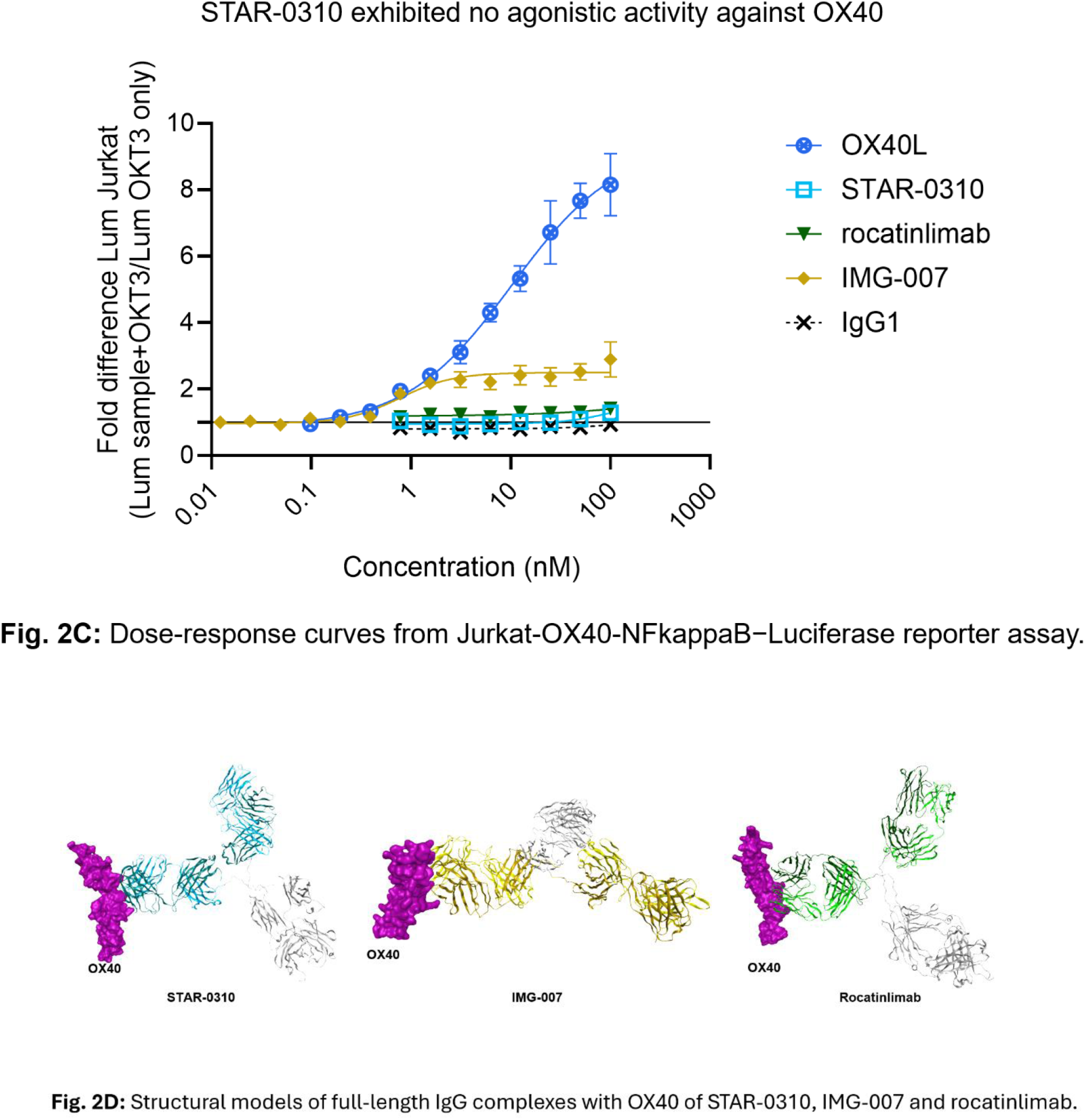
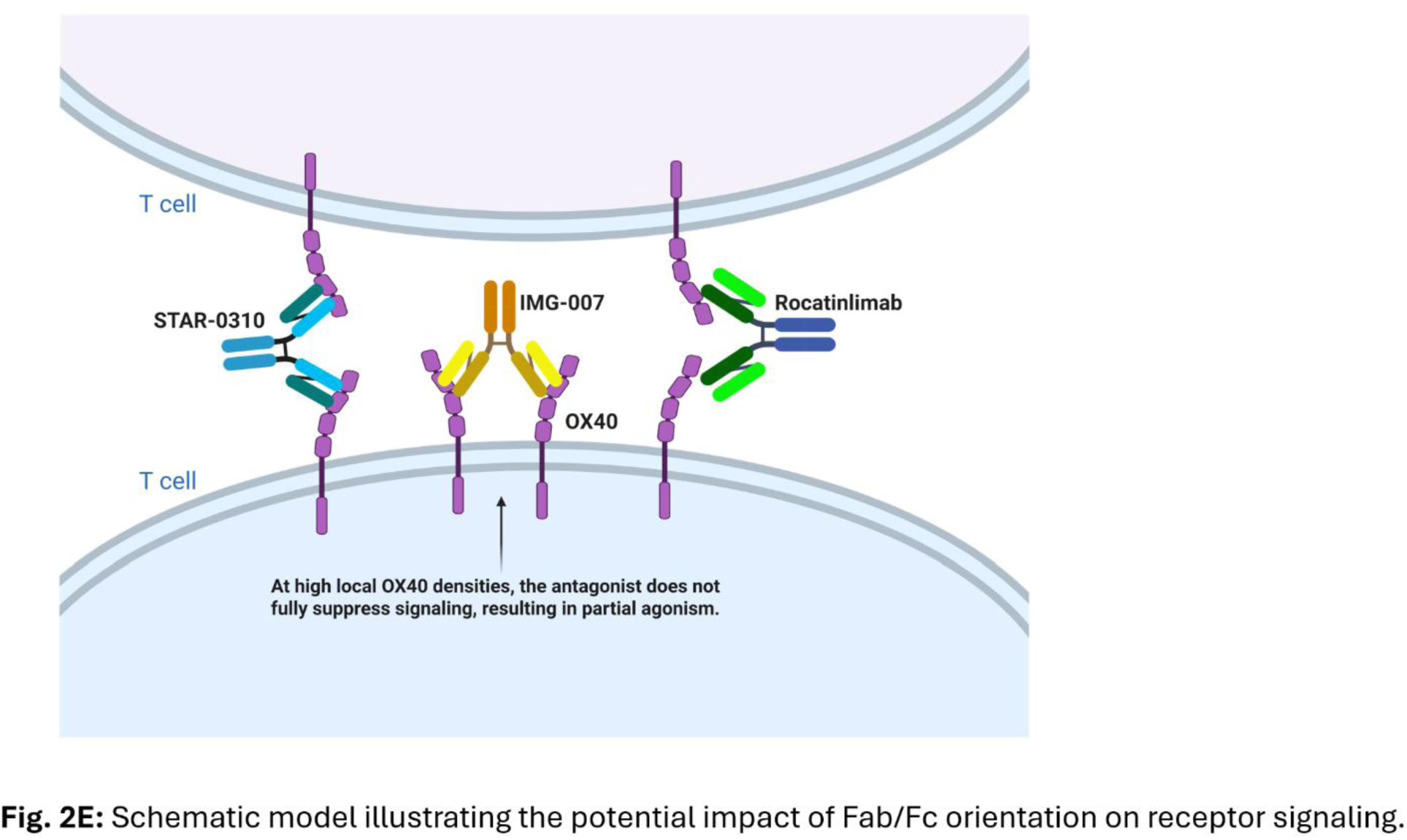
Functional characterization of STAR-0310 demonstrates potent complex disruption and pure antagonism. (A) Surface Plasmon Resonance (SPR) sensorgrams from a complex disruption assay. Biotinylated human OX40L trimer was captured on a sensor chip, followed by injection of human OX40-Fc to form stable complexes. Subsequent injection of antibodies shows that STAR-0310 induces a rapid and substantial dissociation of the pre-formed complexes, at a rate approximately twice that of rocatinlimab and IMG-007. (B) Human OX40 was captured on a sensor chip, followed by injection of biotinylated human OX40L trimer to form OX40/OX40L complex. Subsequent injection of amlitelimab or APG990 analogs shows that the OX40L-targeting antibodies exhibit no disruptive activity. (C) Dose-response curves from a Jurkat-OX40-NFκB−Luciferase reporter assay. The y−axis represents fold−difference in luminescence relative to untreated cells, indicating NF−κB pathway activation. STAR-0310 and rocatinlimab show no increase in luminescence at any concentration, demonstrating pure antagonism similar to the IgG1 isotype control. In contrast, IMG-007 shows a dose-dependent increase in signal, reaching approximately 3-fold activation at higher concentrations, characteristic of a partial agonist. (D) Structural models of full-length IgG complexes with OX40 of STAR-0310, IMG-007 and rocatinlimab depicting the different Fab/Fc orientations. (E) Schematic model illustrating the potential impact of Fab/Fc orientation on receptor signaling. The orientation of STAR-0310 possibly prevents receptor clustering, leading to pure antagonism. In contrast, the distinct orientation of IMG-007 may permit a degree of receptor cross-linking, potentially explaining its partial agonist activity.

A critical characteristic for any antagonist targeting OX40 is the absence of agonistic activity. Since OX40 signaling amplifies T cell proliferation, survival, and cytokine production that drive pathophysiology of AD, any degree of unintended receptor activation by a therapeutic antibody would be counterproductive, potentially limiting its efficacy. Functional activity of STAR-0310 was assessed using a Jurkat-OX40-NFκB-Luciferase reporter assay, which measures OX40-mediated signal transduction. Across all tested concentrations, STAR-0310 exhibited no agonistic activity, behaving as a pure antagonist with a signaling profile comparable to the negative immunoglobulin G (IgG) 1 isotype control and similar to rocatinlimab (Figure 2C). By contrast, IMG-007 induced a dose-dependent, 3-fold increase in NF-κB-driven luminescence compared to untreated controls, indicating partial agonist activity. Structural modeling of the full-length antibodies provides a compelling molecular explanation for this functional divergence (Figures 2D, 2E). The specific Fab and fragment crystallizable (Fc) orientation adopted by STAR-0310 and rocatinlimab upon binding appears to engage OX40 monomers in a conformation that prevents the receptor clustering on the cell surface necessary for signal initiation. Conversely, the distinct binding geometry of IMG-007 may permit a degree of receptor cross-linking, leading to residual signal propagation and the observed partial agonism.

In summary, these analyses demonstrate STAR-0310 possesses a unique dual mechanism of action that distinguishes it from other OX40-targeting therapies. This mechanism comprises: 1) inhibition of the OX40/OX40L interaction via a potent steric hindrance mechanism and 2) efficient disruption of pre-formed, active OX40/OX40L signaling complexes at a rate exceeding that of comparators. Importantly, STAR-0310 functions as a pure OX40 antagonist, exhibiting no detectable intrinsic agonist activity. This differentiated profile suggests that STAR-0310 may achieve comprehensive and durable immunomodulation of T cell-mediated inflammation that drives AD. These conclusions are derived from structural and preclinical studies and have yet to be proved through human clinical trials. These findings provide a strong mechanistic rationale for the ongoing clinical evaluation of STAR-0310 (NCT06782477) as a potential best-in-class therapy for patients with AD.

## Ethics Statement

The Jurkat-OX40-NFkB-Luc reporter assay is an in vitro study using a commercially available cell line and does not require Institutional Review Board approval.

## Data Availability Statement

The data that support the findings of this study are available from the corresponding author upon reasonable request. The Cryo-EM structural data for the STAR-0310/OX40, rocatinlimab/OX40 and IMG-007/OX40 complexes will be deposited in the Electron Microscopy Data Bank (EMDB) and Protein Data Bank (PDB) upon manuscript acceptance.

## Author Contributions

Wrote or contributed to the writing and data interpretation of the manuscript: NB, CZ, JM, JRF, AB, RC, CGB

Participated in research design: NB, CZ, JM

Conducted experiments: NB, CZ, JM, JRF

Performed data analysis: NB, JRF, CZ, JM

## Acknowledgements

Editorial assistance was provided by Jessica Best, DHSc, PA; Michele Gunsior, PhD; Rachel Kilian, PhD; Oluwatobiloba Osikoya, PhD, of Astria Therapeutics. Created in BioRender. Biris, N. (2025) https://BioRender.com/ofnruhg. The authors had full control of the content and made the final decision to submit the manuscript for publication.

## Materials and Methods

### OX40 expression and purification

Human OX40 extracellular domain comprised of residues L29–D170 (OX40 ECD) was cloned into pcDNA3.1 vector containing a C-terminal Avi-His_6_ tag (GLNDIFEAQKIEWHEHHHHHH) and was subsequently amplified and purified using GenElute HP Plasmid Maxiprep Kit (Sigma Aldrich). OX40 ECD was transiently expressed in Expi293F cells (ThermoFisher, Waltham, MA, USA) using ExpiFectamine 293 transfection reagent kit (ThermoFisher). The OX40 ECD was purified from cleared culture supernatant by using HisTrap Excel column (Cytiva, Marlborough, MA, USA) equilibrated with PBS pH 7.4 at 1 mL/min using an Akta Pure 25 FPLC system (Cytiva). Peak fractions are pooled and concentrated using a Centricon centrifugal filter device (Merck Millipore, Burlington, MA, USA) equipped with a 10 kDa cut-off membrane. The concentrated sample was further purified by size exclusion chromatography (SEC) using a HiLoad Superdex-200 16/600 column (Cytiva) in PBS pH 7.4.

The OX40 ECD was subsequently deglycosylated to remove N-linked oligosaccharides and was treated with Protein Deglycosylation Mix II (New England Biolabs, Ipswich, MA, USA), according to manufacturer’s protocol. Briefly, OX40 protein was treated with 1:200 (w/v) Protein Deglycosylation Mix II and was incubated for 18 hours at room temperature.

### Expression and Purification of STAR-0310 and rocatinlimab, IMG-007, amlitelimab and APG990 analogs

For the expression of all antibodies, cDNAs encoding the heavy chain and light chain regions were synthesized and cloned into a pcDNA3.1 derived vector (Invitrogen, Waltham, MA, USA). Publicly available sequences for rocatinlimab, IMG-007, amlitelimab and APG990 analogs were used. The antibodies were transiently expressed in ExpiCHO cells (ThermoFisher) using ExpiFectamine CHO transfection kit (ThermoFisher). Specifically, for the rocatinlimab analog, the culture was supplemented with 100 µM of 2-fluorofucose (SGN-2FF), a fucosyltransferase inhibitor, for the generation of the afucosylated antibody. All antibodies were purified from cleared culture supernatant using HiTrap MabSelect Sure Protein A column (Cytiva) using an Akta Pure 25 FPLC system (Cytiva) followed by SEC using a HiLoad Superdex-200 16/600 (Cytiva) column in PBS pH 7.4.

### Expression and Purification of Fab regions

For the expression of STAR-0310 Fab, cDNAs encoding the heavy chain and light chain regions were synthesized and cloned into a pcDNA3.1 derived vector (Invitrogen).

Equal quantities of heavy chain and light chain vectors were co-transfected into Expi293F cells (ThermoFisher). The expression was then performed as described previously for human OX40 ECD. After 5 days post-transfection, cell-free culture supernatant containing the recombinant Fab region was prepared by centrifugation followed by filtration and used for further purification. The supernatant was loaded on a CaptureSelect CH1-XL column (Thermo Fisher Scientific) equilibrated with PBS pH 7.4. The Fab fragment was then eluted with 0.1 M glycine, pH 3.0. After neutralization with 1/20 volume of Tris-HCl pH 8.0, the eluate was further purified by SEC using a HiLoad Superdex-200 16/600 column (Cytiva) equilibrated in PBS pH 7.4.

The Fab fragments of rocatinlimab and IMG-007 were obtained by papain digestion using the Pierce Fab preparation kit (Thermo Fisher Scientific) according to manufacturer’s instructions. Briefly, IgG samples were buffer exchanged into digestion buffer at pH 7.0 using Zeba Spin Desalting columns. Immobilized papain resin was equilibrated with digestion buffer and incubated with the IgG sample at 37°C. Digestion was terminated by centrifugation to separate the Fab-containing supernatant from the resin. Fab fragments were purified by passing the digest through a NAb Protein A Plus Spin Column, which retained Fc fragments and undigested IgG.

### Cryo-EM Sample Preparation and Data Acquisition

For application to cryo-EM grids, OX40:Fab complexes were adjusted to 2.25 – 2.5 mg/mL concentration and supplemented with 0.125% (w/v) CHAPSO detergent. Sample concentration, grid type, and microscopy details are given in Supplemental Figure X. Grids were glow discharged immediately prior to use. Sample grid blotting and plunge freezing was performed using a Vitrobot Mk IV (Thermo Fisher Scientific) with the blotting chamber held at 100% humidity and 18 °C. A 3.5-µl droplet of sample was applied to the grid, blotted for 4 s and then plunged into liquid ethane. Cryo-EM imaging was performed using Krios G3i electron microscopes (Thermo Fisher Scientific) equipped with BioQuantum K3 energy filters and cameras (Gatan, Inc, Pleasanton, CA, USA). Acquisition details are given in Supplemental Table 1.

### Cryo-EM Data Processing

All cryo-EM data processing was carried out using RELION 5.0^1^. A summary of the datasets and final map refinement results is given in Supplemental Table 2. Exposure movies were motion corrected using the patch-based implementation in RELION and micrograph contrast transfer functions (CTF) were estimated using CTFFind 4.1.14^2^ from movie amplitude spectra sums. Exposures that failed- or were outliers in-fitted motion or CTF estimation parameters, and micrographs with significant non-vitreous ice contamination, were removed from downstream processing. For the STAR-0310 and datasets, particles were selected from the entire micrograph set first using Topaz^3^ based on its default/general pre-trained model. These picks were processed through multiple rounds of 2D classification and 3D initial model/3D classification to generate a 3D reference map which was used for reference-based picking of the entire micrograph set. Reference-based picks were similarly filtered through 2D and 3D classification and pooled with the Topaz picks, removing overlapping/duplicate particles. For the IMG-007 analog and rocatinlimab analog datasets, a random subset of 1000 micrographs was subjected to the process described above. The resulting particle positions were used to train a Topaz model that was then used to pick the entire micrograph set, which were again filtered through multiple rounds of 2D and 3D classification.

For all datasets, the surviving particles were then cycled through CTF refinement, per-particle motion correction (Bayesian polishing), and 3D classification without particle pose searches (with 3D auto-refinement between each step) until global map resolution ceased to improve. All 3D auto-refinement and 3D classification steps utilized a soft solvent mask, and all 3D auto-refinement steps utilized Blush^4^ (STAR-0310) or SIDESPLITTER^5^ (IMG-007 and rocatinlimab) for regularization.

### Cryo-EM Model Building

Initial atomic models were generated using ModelAngelo^6^. These models were then further built and refined by iterating between manual adjustments in Coot^7^ and automated real-space refinement in Phenix^8^. Manual modeling in Coot utilized a combination of maps that had been (1) globally sharpened by an automatically determined B-factor and filtered to local resolution (using the procedure implemented in RELION), and (2) sharpened and filtered by LAFTER^9^. Automated refinement and all map/model validation in Phenix utilized the map output from RELION 3D auto-refinement without any modification. Model geometry and map-model fit statistics are given in Supplemental Table 3.

### Surface Plasmon Resonance displacement assay

Surface Plasmon Resonance (SPR) analysis was used to determine the ability of STAR-0310, rocatinlimab and IMG-007 antibodies to disrupt the preformed OX40L-OX40 complex.

Measurements were performed on a Biacore 8K+ instrument (Cytiva Life Sciences) using the Biacore 8K+ Control Software and analyzed with the Biacore Insight Evaluation Software (v3.0). Briefly, an anti-Avi tag antibody (Genscript) was immobilized at around ≈ 7,000-9,000 resonance units (RU) on a Series S CM5 Sensor Chip (Cytiva Life Sciences) using an amine coupling kit (Cytiva Life Sciences) in both flow cells. Biotinylated human OX40L-His-Avi (Acro Biosystems) protein was injected into flow cell 2 at 50 nM followed by human OX40-Fc (Acro Biosystems) protein being injected into both flow cells at 200 nM. The evaluation was performed by injecting STAR-0310, rocatinlimab, and IMG-007 at 1340 nM in both flow cells, in triplicates. After each binding event, baseline levels were restored using a regeneration solution (75 mM phosphoric acid) on both flow paths. HBS-EP+ pH 7.4 solution (Cytiva Life Sciences) was used as a running buffer. Data was analyzed using the Multi-Cycle Kinetics (MCK) with capture evaluation method of the Biacore Insight Evaluation Software and off-rate values for OX40-OX40L dissociation were determined using the 1:1 dissociation fitting model.

The ability of the OX40L-targeted antibodies amlitelimab and APG990 to disrupt the preformed OX40L-OX40 complex was assessed using an SPR displacement assay. Briefly, human OX40-His protein (Acro Biosystems) was immobilized at about 800-1000 RU on a CM5 Series S sensor chip (Cytiva Life Sciences) using an amine coupling kit (Cytiva Life Sciences) in both flow cells. Biotinylated human OX40L-His-Avi (Acro Biosystems) protein was injected in both flow cells at 100 nM for 120 sec.

Subsequently, Amlitelimab and APG990 analogs were injected for 150 sec at 1340 nM concentration in both flow cells, in triplicates.

### Jurkat-OX40-NFκB-luciferase reporter agonism assay

Jurkat-NF-κB-OX40 or OX40 Effector Cells, a genetically engineered Jurkat T cell line that expresses human OX40 and an NF-κB luciferase reporter (Promega), were used to assess the potential agonistic activity of STAR-0310 in comparison to comparator antibodies rocatinlimab, IMG-007, and a control antibody. The aim of this assay was to measure the ability of the tested molecules to induce agonism through the activation of the NFĸB pathway in Jurkat-NFĸB-OX40 reporter cells. In brief, 0.1×10^6^ cells/well Jurkat cells were plated in sterile 96-well µCLEAR microplates (VWR) coated with anti-CD3 Monoclonal Antibody OKT3 (Invitrogen, 5 ug/ml) and incubated in the presence of STAR-0310, rocatinlimab, IMG-007 or a negative control (irrelevant binder). Control wells without OKT3 coating were included. Treatments, OX40L positive control and IgG1 isotype negative control were prepared 3x concentrated to reach a final concentration of 100 nM in the wells, followed by a 1/2 serial dilution. The tested molecules and controls were added to the corresponding wells and incubated for 5 hours at 37°C and 5% CO_2_. Maximum luminescence was assessed by coating OKT3 without treatment. This control well is called in the formula as “OKT3 only”. After incubation, 75 µl of Bio-Glo solution reagent (Promega) was added to the wells and incubated for 15 minutes. Luminescence was measured using a Synergy Neo Plate Reader. The luminescence level from OKT3 stimulation was considered the baseline level and the fold difference between the anti-OX40 antibody induced luminescence vs. OKT3 base line luminescence was calculated using the following formula:

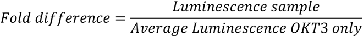

A logistic 4 parameters non-linear regression model with variable slope was applied to all datasets and maximal luminescence and maximal fold difference were determined. The analysis was done with GraphPad (v. 10.4.2). No statistical analysis was performed due to the limited number of repeats (n:2-3 repeats) for all molecules.

---

**Supplemental Table 1.**
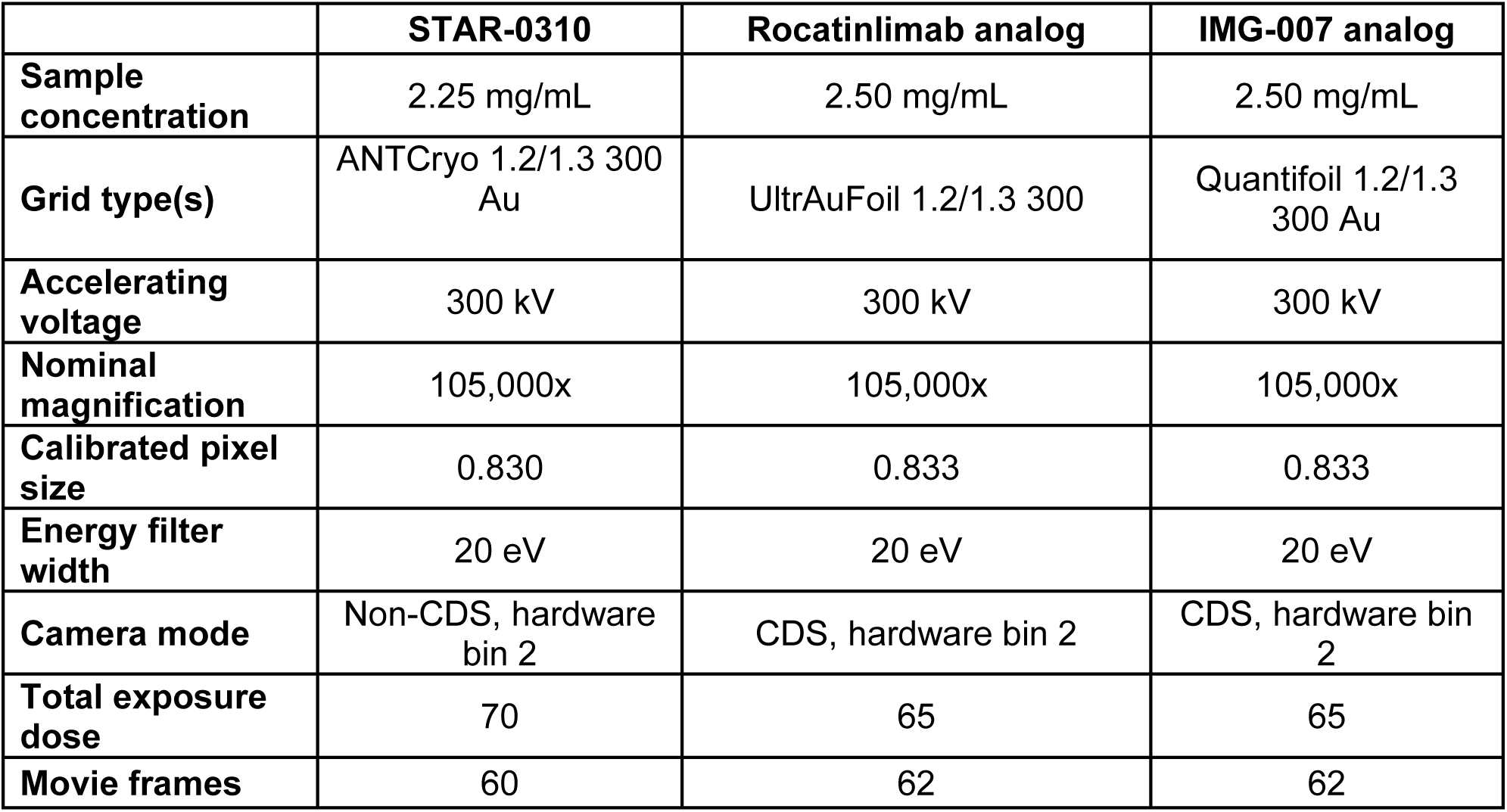
Acquisition details for the STAR-0310, rocatinlimab and IMG-007 analogs.

**Supplemental Table 2.**
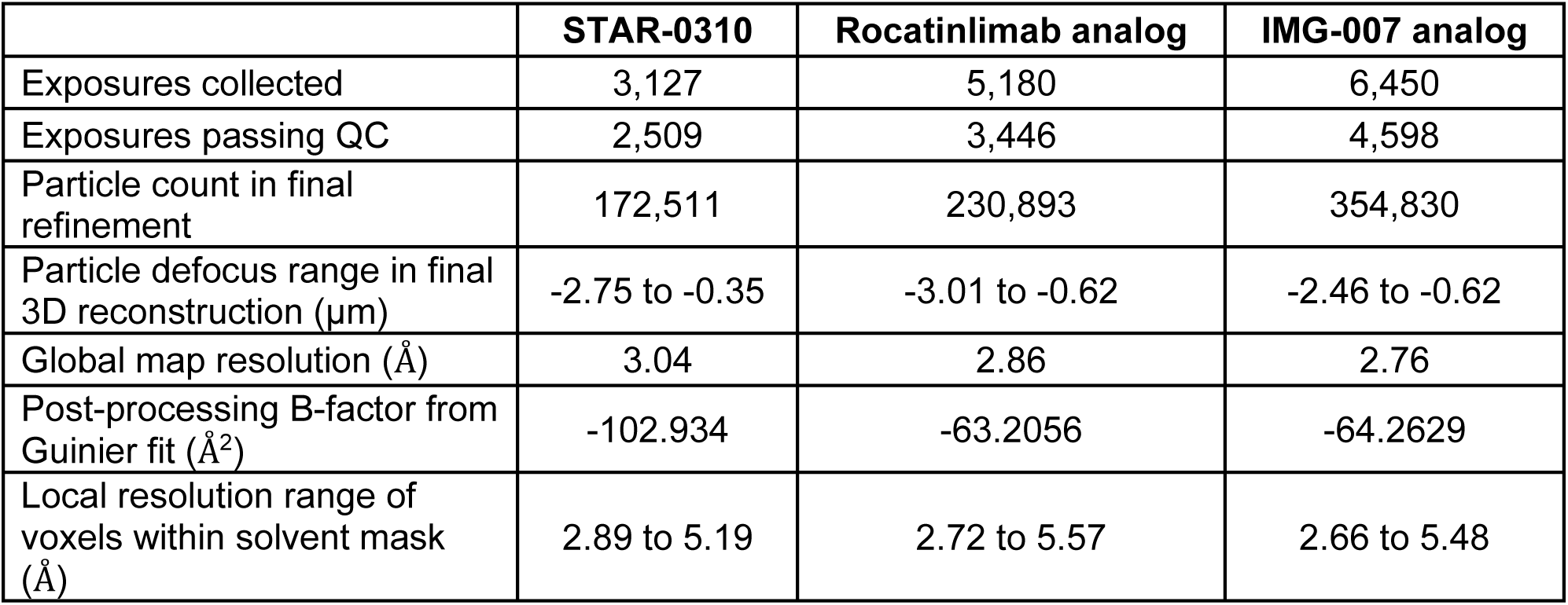
Summary of the datasets and final map refinement results.

**Supplemental Table 3.**
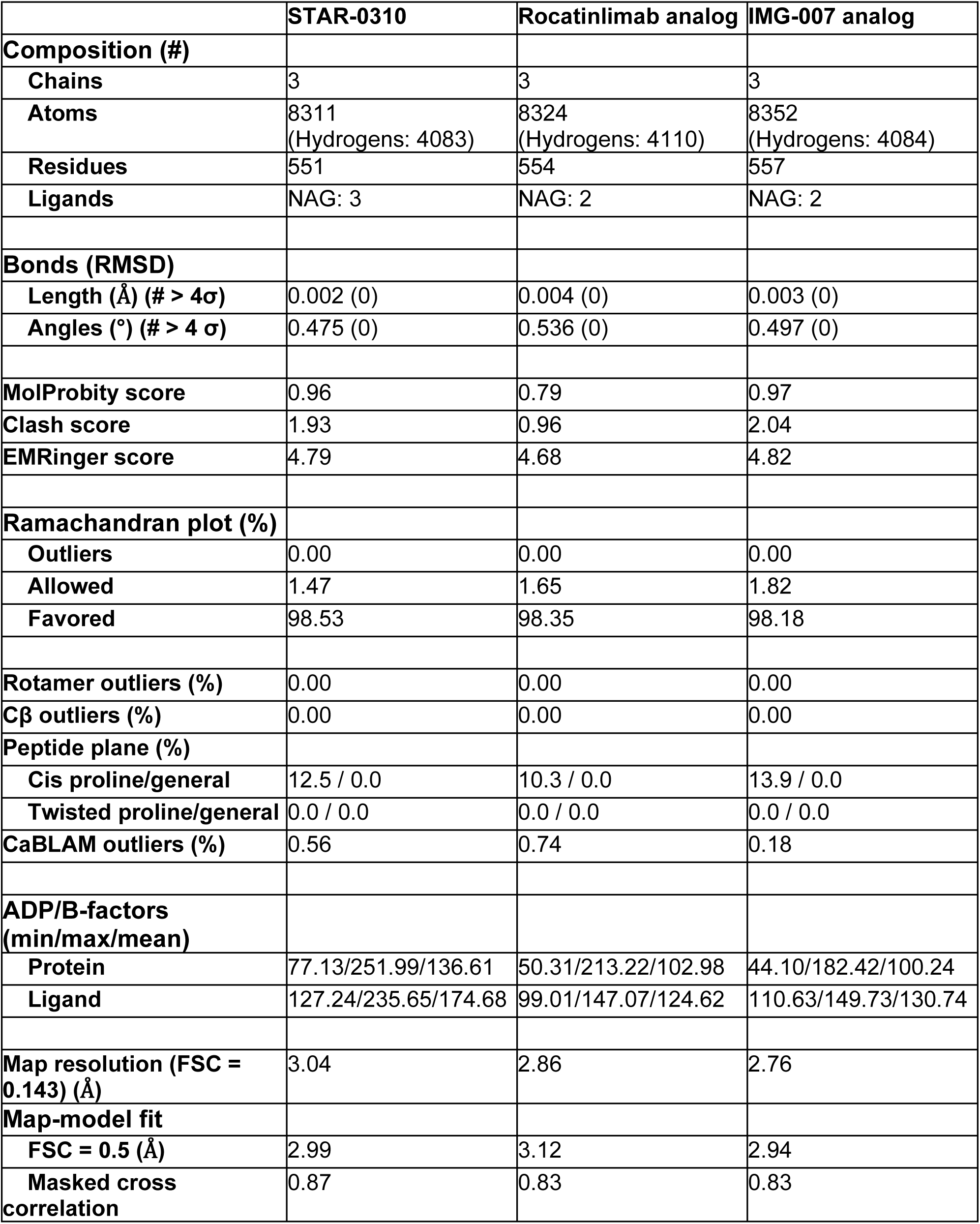
Model geometry and map-model fit statistics.

