## Supplementary Materials for "A Dual-Action Mechanism to Prevent OX40 Signaling: The Structural Basis for the Differentiated Antagonist STAR-0310"

**Supplemental Table 1.** Acquisition details for the STAR-0310, rocatinlimab and IMG-007 analogs.

| | STAR-0310 | Rocatinlimab analog | IMG-007 analog |
|---|---|---|---|
| **Sample concentration** | 2.25 mg/mL | 2.50 mg/mL | 2.50 mg/mL |
| **Grid type(s)** | ANTCryo 1.2/1.3 300 Au | UltrAuFoil 1.2/1.3 300 | Quantifoil 1.2/1.3 300 Au |
| **Accelerating voltage** | 300 kV | 300 kV | 300 kV |
| **Nominal magnification** | 105,000x | 105,000x | 105,000x |
| **Calibrated pixel size** | 0.830 | 0.833 | 0.833 |
| **Energy filter width** | 20 eV | 20 eV | 20 eV |
| **Camera mode** | Non-CDS, hardware bin 2 | CDS, hardware bin 2 | CDS, hardware bin 2 |
| **Total exposure dose** | 70 | 65 | 65 |
| **Movie frames** | 60 | 62 | 62 |

**Supplemental Table 2.** Summary of the datasets and final map refinement results.

| | STAR-0310 | Rocatinlimab analog | IMG-007 analog |
|---|---|---|---|
| Exposures collected | 3,127 | 5,180 | 6,450 |
| Exposures passing QC | 2,509 | 3,446 | 4,598 |
| Particle count in final refinement | 172,511 | 230,893 | 354,830 |
| Particle defocus range in final 3D reconstruction (μm) | -2.75 to -0.35 | -3.01 to -0.62 | -2.46 to -0.62 |
| Global map resolution (Å) | 3.04 | 2.86 | 2.76 |
| Post-processing B-factor from Guinier fit (Å$^2$) | -102.934 | -63.2056 | -64.2629 |
| Local resolution range of voxels within solvent mask (Å) | 2.89 to 5.19 | 2.72 to 5.57 | 2.66 to 5.48 |

**Supplemental Table 3.** Model geometry and map-model fit statistics.

| | STAR-0310 | Rocatinlimab analog | IMG-007 analog |
|---|---|---|---|
| **Composition (#)** | | | |
| **Chains** | 3 | 3 | 3 |
| **Atoms** | 8311 (Hydrogens: 4083) | 8324 (Hydrogens: 4110) | 8352 (Hydrogens: 4084) |
| **Residues** | 551 | 554 | 557 |
| **Ligands** | NAG: 3 | NAG: 2 | NAG: 2 |
| **Bonds (RMSD)** | | | |
| **Length (Å) (# > 4σ)** | 0.002 (0) | 0.004 (0) | 0.003 (0) |
| **Angles (°) (# > 4 σ)** | 0.475 (0) | 0.536 (0) | 0.497 (0) |
| **MolProbity score** | 0.96 | 0.79 | 0.97 |
| **Clash score** | 1.93 | 0.96 | 2.04 |
| **EMRinger score** | 4.79 | 4.68 | 4.82 |
| **Ramachandran plot (%)** | | | |
| **Outliers** | 0.00 | 0.00 | 0.00 |
| **Allowed** | 1.47 | 1.65 | 1.82 |
| **Favored** | 98.53 | 98.35 | 98.18 |
| **Rotamer outliers (%)** | 0.00 | 0.00 | 0.00 |
| **Cβ outliers (%)** | 0.00 | 0.00 | 0.00 |
| **Peptide plane (%)** | | | |
| **Cis proline/general** | 12.5 / 0.0 | 10.3 / 0.0 | 13.9 / 0.0 |
| **Twisted proline/general** | 0.0 / 0.0 | 0.0 / 0.0 | 0.0 / 0.0 |
| **CaBLAM outliers (%)** | 0.56 | 0.74 | 0.18 |
| **ADP/B-factors (min/max/mean)** | | | |
| **Protein** | 77.13/251.99/136.61 | 50.31/213.22/102.98 | 44.10/182.42/100.24 |
| **Ligand** | 127.24/235.65/174.68 | 99.01/147.07/124.62 | 110.63/149.73/130.74 |
| **Map resolution (FSC = 0.143) (Å)** | 3.04 | 2.86 | 2.76 |
| **Map-model fit** | | | |
| **FSC = 0.5 (Å)** | 2.99 | 3.12 | 2.94 |
| **Masked cross correlation** | 0.87 | 0.83 | 0.83 |
